# Localization of planteose hydrolysis during seed germination of *Orobanche minor*

**DOI:** 10.1101/2021.06.16.448768

**Authors:** Atsushi Okazawa, Atsuya Baba, Hikaru Okano, Tomoya Tokunaga, Tsubasa Nakaue, Takumi Ogawa, Shuichi Shimma, Yukihiro Sugimoto, Daisaku Ohta

## Abstract

Root parasitic weeds of the Orobanchaceae, such as witchweeds (*Striga* spp.) and broomrapes (*Orobanche* and *Phelipanche* spp.), cause serious losses in agriculture worldwide. No practical method to control these parasitic weeds has been developed to date. Understanding the characteristic physiological processes in the life cycles of root parasitic weeds is particularly important to identify specific targets for growth modulators. In our previous study, planteose metabolism was revealed to be activated soon after the perception of strigolactones in germinating seeds of *O*. *minor*. Nojirimycin inhibited planteose metabolism and impeded seed germination of *O*. *minor*, indicating that planteose metabolism is a possible target for root parasitic weed control. In the present study, we investigated the distribution of planteose in dry seeds of *O*. *minor* by matrix-assisted laser desorption/ionization–mass spectrometry imaging. Planteose was detected in tissues surrounding—but not within—the embryo, supporting its suggested role as a storage carbohydrate. Biochemical assays and molecular characterization of an α-galactosidase family member, OmAGAL2, indicated the enzyme is involved in planteose hydrolysis in the apoplast around the embryo after the perception of strigolactones to provide the embryo with essential hexoses for germination. These results indicated that OmAGAL2 is a potential molecular target for root parasitic weed control.

**Highlight:** Planteose accumulated in tissues surrounding the embryo in *Orobanche minor* dry seeds and was indicated to be hydrolyzed in the apoplast around the embryo by α-galactosidase during germination.

## Introduction

Root parasitic weeds belonging to the Orobanchaceae, such as witchweeds (*Striga* spp.) and broomrapes (*Orobanche* and *Phelipanche* spp.), cause serious losses in agriculture worldwide (Parker, 2013). Many studies seeking to identify an effective solution have been conducted (Fernández-Aparicio *et al.*, 2020). However, no practical method to control theses parasitic weeds has been developed to date. One potential method is utilization of growth modulators specific to the root parasitic weeds. Therefore, understanding the characteristic physiological processes in the life cycles of root parasitic weeds is particularly important to identify specific targets of growth modulators.

The life cycle of root parasitic weeds is dependent on their hosts, because the weeds are obligate parasites. Germination of their seeds is host-dependent and requires germination stimulants, in most cases, strigolactones (SLs) (Brun *et al.*, 2018; Bouwmeester *et al.*, 2019, 2021; Yoneyama, 2020). Strigolactones are released from plant roots as signals to establish symbiotic relationships with arbuscular mycorrhizal fungi (Akiyama *et al.*, 2005; López-Ráez, 2015; Lanfranco *et al.*, 2018). Strigolactones function *in planta* as a hormone that alters plant architecture in response to the nutritional status (Gomez-Roldan *et al.*, 2008; Umehara *et al.*, 2008; Al-Babili and Bouwmeester, 2015; Waters *et al.*, 2017; Bürger and Chory, 2020). Recent biochemical studies revealed that the mode of perception of SLs as a germination stimulant in root parasitic weeds is different from that as a plant hormone (Tsuchiya *et al.*, 2015, Khosla and Nelson, 2016). In Arabidopsis (*Arabidopsis thaliana*) and rice (*Oryza sativa*), SLs are perceived by the SL receptor DWARF14/DECREASED APICAL DOMINANCE2 (D14/DAD2) (Arite *et al.*, 2009; Hamiaux *et al.*, 2012; Nakamura *et al.*, 2013; Seto *et al.*, 2019). However, in *Striga hermonthica*, SLs are perceived by a subset of KARRIKIN INSENSITIVE2 (KAI2)/HYPOSENSITIVE TO LIGHT (HTL) receptors (KAI2ds) (Conn *et al.*, 2015; Tsuchiya *et al.*, 2015; Conn and Nelson, 2016;). *KAI2d* genes are paralogous to *D14* and are highly diversified in the Orobanchaceae (Conn *et al.*, 2015; Tsuchiya *et al.*, 2015; Conn and Nelson, 2016; Yoshida *et al.*, 2019). The diversification might be closely linked to SL-dependent germination and the host specificity of the parasitic species (Khosla and Nelson, 2016). Precise characterization of the binding affinity of KAI2d/HTLs in *S*. *hermonthica* with SLs have led to selection and design of a prominent SL agonist, sphynolactone-7 (SPL-7) (Uraguchi *et al.*, 2018).

In comparison with SL perception in root parasitic weeds, metabolic processes in germinating seeds after SL perception are poorly understood. Previously, we conducted metabolome analyses on germinating seeds of *Orobanche minor* and revealed that metabolism of planteose (α-D-galactopyranosyl-(1→6)-β-D-fructofuranosyl-(2→1)-α-D-glucopyranoside) is activated soon after the perception of a synthetic SL, *rac*-GR24 (Wakabayashi *et al.*, 2015). We also observed that nojirimycin (NJ) inhibits planteose metabolism and impedes the germination of *O*. *minor* seeds (Wakabayashi *et al.*, 2015, Harada *et al.*, 2017). Transcriptome analysis of the effect of NJ on *O*. *minor* germination suggested that NJ inhibits planteose metabolism through disturbance of sugar signaling (Okazawa *et al.*, 2020). These studies indicate that planteose metabolism and the subsequent change in sugar profile in the seeds are critical processes in the germination of *O*. *minor*.

Planteose is accumulated in the seeds of certain plant species, such as, tobacco (*Nicotiana tabacum*) (French, 1955), mints (Lamiaceae spp.) (French *et al.*, 1959), sesame (*Sesamum indicum*) (Dey, 1980), and chia (*Salvia hispanica*) (Xing *et al.*, 2017). In addition to *O*. *minor*, seeds of the Orobanchaceae root parasitic weeds *O*. *crenata*, *P*. *aegyptiaca*, and *S*. *hermonthica* contain planteose and its hydrolysis is rapidly induced after perception of *rac*-GR24 (Wakabayashi *et al.*, 2015). We previously revealed that the first step of planteose metabolism is hydrolysis of the α-galactosyl linkage to produce Suc and Gal (Wakabayashi *et al.*, 2015). However, the enzymes involved in the metabolism and biosynthesis of planteose in plants remain to be elucidated.

α-Galactosidases (EC 3.2.1.22; AGALs) in plants are characterized in relation to metabolism of raffinose family oligosaccharides (RFOs) (Van den Ende, 2013) and galactomannan (Buckeridge *et al.*, 2000) during seed germination. Acid AGALs belong to the plant α-galactosidase subfamily in family 27 of glycoside hydrolases (GH27) (Van den Ende, 2013; Imaizumi *et a*l., 2017), whereas alkaline AGALs belong to GH36 (Van den Ende, 2013). Seeds of various plant species contain RFOs (Kuo *et al.*, 1988), which are hydrolyzed by AGALs during germination. In pea (*Pisum sativum*), constant activity of acid AGAL was confirmed during germination, whereas alkaline AGAL was expressed after radicle protrusion (Blöchl *et al.*, 2008). Increase in acid AGAL activity to hydrolyze RFOs or galactomannan has been confirmed in soybean (*Glycine max*) (Guimarães *et al.*, 2001) and *Tachigali multijuga* (members of the Fabaceae) (Fialho *et al.*, 2008). Acid AGAL activity also increases in the micropylar region of the endosperm of tomato (*Solanum lycopersicum*) (Feurtado *et al.*, 2001) and date palm (*Phoenix dactylifera*) (Chandra Sekhar and DeMason, 1990) during germination. Transcripts of a gene encoding AGAL were isolated from aleurone cells in germinating seeds of guar (*Cyampsis tetragonaloba*), suggesting that AGAL is secreted into the endosperm for degradation of storage galactomannan (Hughes *et al.*, 1988). These results indicate that AGALs are involved in the metabolism of storage carbohydrates in the endosperm to promote seed germination in a number of plant species.

In this study, we investigated AGAL activities during seed germination of *O. minor* together with planteose distribution in the dry seeds, to gain insight into the metabolism driving germination after perception of germination stimulants. Cloning and characterization of a member of the acid AGALs subfamily, OmAGAL2, indicated its involvement in planteose hydrolysis. The present results suggested that planteose is stored outside of the embryo in the dry seeds, after SL perception, is hydrolyzed by OmAGAL2 in the apoplast to supply Suc, and subsequentially released hexoses, to the embryo.

## Materials and methods

### Plant material and germination treatment

Seeds from wild-grown *Orobanche minor* plants were collected in Yokohama, Japan in June, 2013. Seed germination was induced as described previously (Wakabayashi *et al.*, 2015). The seeds were surface-sterilized with 1% sodium hypochlorite containing 0.1% (w/v) Tween 20 for 2 min at 42 °C with shaking, then rinsed several times with distilled water, and dried under vacuum. The dried seeds were placed on two layers of glass microfiber filters (Whatman GF/D, GE Healthcare, Chicago, IL, USA) moistened with distilled water in a Petri dish in the dark at 23 °C for a week (conditioning). After conditioning, the seeds on the upper layer of the glass filter were transferred to a new Petri dish with a single glass microfiber filter, and strigolactone solution (*rac*-GR24, final concentration 1 mg L^−1^) was applied. To visualize α-galactosidase activity *in vivo*, 5-bromo-4-chloro-3-indolyl-α-D-galactopyranoside (X-α-Gal; 40 µg mL^−1^) was applied directly to the germinating seeds.

### Purification of planteose from chia seeds

Planteose was extracted in accordance with the method reported previously (Xing *et al.*, 2017) with minor modifications. Dry chia (*Salvia hispanica*) seeds (1 g) were incubated in 20 mL distilled water at 80 °C for 2 h. A viscous fluid was obtained by passing the liquid through a drain net, and was then frozen in liquid nitrogen and freeze-dried. Aqueous EtOH (70%) was added to the dried sample, stirred for 10 min, and centrifuged at 4,000 *g* for 10 min. Planteose was purified by HPLC comprising a system controller (SCL-10Avp, Shimadzu, Kyoto, Japan), a column oven (CTO-10A, Shimadzu), a pump (LC-10AT, Shimadzu), and a refractive index detector (RID-10A, Shimadzu) equipped with a COSMOSIL Sugar-D packed column (20 mm ID × 250 mm, Nacalai Tesque, Kyoto, Japan) at 35 °C. A mixture of acetonitrile–water (65:35, v/v) was used as the mobile phase at the flow rate of 1 mL min^−1^ with an isocratic mode.

### MALDI–MSI

2,5-Dihydroxybenzoic acid (2,5-DHB) and ultrapure water were purchased from Merck (Darmstadt, Germany). Indium-thin-oxide (ITO) glass slides (SI0100N) for matrix-assisted laser desorption/ionization–mass spectrometry imaging (MALDI–MSI) analysis were purchased from Matsunami Glass (Osaka, Japan). As an embedment medium, 4% carboxymethyl cellulose (CMC) was purchased from Section Lab (Yokohama, Japan). The 2,5-DHB solution was prepared at the concentration of 50 mg mL^−1^ using 50% MeOH. Planteose solution (0.01, 0.1, or 1.0 mg mL^−1^ in ultrapure water) was mixed with 2,5-DHB solution of the same concentration and the mixed solution was spotted onto the ITO glass slide. After crystallization, the spots were analyzed to optimize laser power and detector voltage.

Seeds of *O*. *minor* (1 mg) and 500 µL of 4% CMC were mixed, and then the mixture was placed in a disposable base mold (Tissue-Tek, Sakura Finetek, Tokyo, Japan). After freezing at −80 °C, tissue sectioning was performed using a cryomicrotome (CM1950, Leica, Nussloch, Germany). The obtained tissue sections were thaw-mounted onto ITO glass slides. The glass slides were air-dried and then subjected to matrix application. The matrix, 2,5-DHB, was heated at 180 °C and sublimated at a thickness of 1.0 µm using a vacuum deposition system (iMLayer, Shimadzu). After formation of the matrix layer on the sample surface, the imaging experiment was performed using an iMScope TRIO (Shimadzu). Based on the results of standard sample analysis, the target peak was identified at *m/z* 527.16. To improve the specificity of the imaging results, MS/MS imaging was performed. The peak at *m/z* 527.16 was isolated inside the ion trap and dissociation was induced using argon gas. The target fragment peaks were *m/z* 347.11 and *m/z* 365.11. To confirm the accuracy of the imaging result, both peak intensity maps were generated. In the imaging experiment, the laser diameter was 5 µm and the interval of data points was 3 µm.

### Protein extraction from germinating seeds of O. minor

Protein extraction was conducted as described previously (Wakabayashi *et al.*, 2015) at 4 °C with minor modifications. Germinating seeds (ca. 40 mg) of *O*. *minor* were frozen in liquid nitrogen and disrupted with a ball mill (20 Hz, 5 min; MN301, Verder Scientific, Haan, Germany). Extraction buffer (1.5 mL, pH 7.0) composed of 50 mM HEPES, 1 mM DTT, 1 mM EDTA, 20 mg polyvinylpolypyrrolidone (PVPP), and 1 % Protease Inhibitor Cocktail (Merck) was added and incubated for 5 min. The homogenate was centrifuged at 12,000 *g* for 15 min and 0.8 mL of supernatant was collected. The residue was re-extracted in extraction buffer without PVPP, and 0.7 mL of extract was collected after centrifugation and combined with the first extract. The combined extracts (1.5 mL) was used as a soluble enzyme fraction. The enzyme extract was desalted on a PD-10 column (Cytiva, Marlborough, MA, USA) previously equilibrated with 50 mM HEPES buffer (pH 7.0) and concentrated with an Amicon Ultra-4 10K centrifugal filter (Merck).

### α-Galactosidase assay

*p*-Nitrophenyl-α-D-galactopyranoside (*p*-NP-α-Gal) (Tokyo Chemical Industry, Tokyo, Japan) was used as a model substrate to measure α-galactosidase activity. Proteins (5 µg) were incubated with the substrate (final concentration 1.0 mM) in 0.1 M citrate buffer (pH 3.0−6.0) or 0.1 M phosphate buffer (pH 6−8) at 37 °C for 30 min, and then 0.5 M Na_2_CO_3_ was added to stop the reaction. The amount of released *p*-nitrophenol was quantified by measuring absorbance at 410 nm with a microplate leader (SH-9000, Corona Electric, Hitachinaka, Japan). When planteose was used as a substrate, the reaction was monitored using an ultra-performance liquid chromatograph equipped with an evaporative light scattering detector (UPLC–ELSD; ACQUITY, Waters, Milford, MA, USA) as reported previously (Wakabayashi *et al*, 2015). Sugars were separated by an ACQUITY UPLC BEH Amide 1.7 µm column (2.1 mm ID × 150 mm, Waters) at 35 °C. The mobile phase was acetonitrile with 0.2% triethylamine (TEA; solvent A) and ultrapure water with 0.2% TEA (solvent B). Separation was performed using a linear gradient program as follows: 20–30% B for 0–2.8 min, 30–50% B for 2.8–4.5 min, 50–80% B for 4.5–5.0 min, 80–20% B for 5.0–6.5 min, and 20% B for 6.5–7.0 min. The flow rate was set as follows: 0.25 mL min^−1^ for 0–4.5 min, 0.25–0.10 mL min^−1^ for 4.5–5.0 min, 0.10–0.25 mL min^−1^ for 5.0–6.5 min, and 0.25 mL min^−1^ for 6.5–7.0 min. The sample injection volume was 5 µL.

### RT-qPCR

Total RNA was isolated from germinating seeds of *O*. *minor* using TRIzol Reagent (Thermo Fischer Scientific, Waltham, MA, USA) and treated with the RNase-Free DNase Set (Qiagen, Hilden, Germany) in accordance with the manufacturer’s instructions. Synthesis of cDNA and quantitative PCR was conducted using the PrimeScript RT Master Mix (Takara Bio, Kusatsu, Japan) with Thermal Cycle Dice TP800 (Takara Bio). The sequences of gene-specific primers used were as follows: *OmAGAL2* Fw, 5′-GGGATGACTGCCGAAGAATA-3′; *OmAGAL2* Rv, 5′-TGCTTAGTGTCGCCAATGTC-3′; comp71446_c0_seq1 Fw, 5′-TGGATTTGCTGGAGTTGGTG-3′; and comp71446_c0_seq1 Rv, 5′-CGATGGGAATTCAGACGACA-3′. The gene comp71446_c0_seq1 was shown to be constitutively expressed in our previous transcriptome analysis (Okazawa *et al*, 2020), and was selected as a reference gene for normalization using the 2^−ΔCt^ method.

### Expression of OmAGAL2 in Escherichia coli

Genes encoding α-galactosidases (*OmAGALs*) were surveyed in the transcriptome in germinating seeds of *O*. *minor* (Okazawa *et al.*, 2020). The *OmAGAL2* coding sequence without the expected signal peptide (SP) sequence (*ΔSP-OmAGAL*, amino acids [AA] 47–412) was cloned into the pENTR/D-TOPO vector (Thermo Fisher Scientific) in accordance with the manufacturer’s instructions, then subcloned into the pGEX-5X-1-GW vector (Okazawa *et al.*, 2014) using the Gateway LR reaction. *Escherichia coli* BL21(DE3) was transformed with the constructed expression vector, pGEX-ΔSP-OmAGAL2. The transformed *E*. *coli* cells were pre-cultured in LB medium supplemented with 100 µg mL^−1^ ampicillin for 18 h at 37 °C. A portion of the pre-culture (1/250) was transferred to new LB medium supplemented with 100 µg mL^−1^ ampicillin and cultured at 37 °C until OD_600_ = 0.45 was attained. Protein expression was induced by addition of isopropyl β-D-thiogalactoside (final concentration 0.1 mM). After incubation at 9 °C for 72 h, the cultured cells were harvested by centrifugation (400 *g*, 4 °C, 15 min). The harvested cells were sonicated with an ultrasonic disruptor (UR-20P, Tomy Seiko, Tokyo, Japan) in buffer (10 mM Na_2_HPO_4_, 1.8 mM KH_2_PO_4_, 137 mM NaCl, 2.7 mM KCl, and 1 mM dithiothreitol, pH 7.4) for 15 s for a total of five times. After centrifugation (400 *g*, 4 °C, 15 min), the supernatant was collected as the crude enzyme solution. Recombinant GST-ΔSP-OmAGAL2 was purified using the GST SpinTrap column (Cytiva) in accordance with the manufacturer’s instruction manual. The glutathione *S*-transferase (GST) tag was cleaved using Factor Xa Protease (New England Biolabs, Ipswich, MA, USA) in accordance with the manufacturer’s instruction.

### Subcellular localization analyses

*OmAGAL2*, *ΔSP-OmAGAL2*, and *SP* (AA1–46) were cloned separatory into the mC121 vector to express proteins fused with mCherry at the C-terminus under the control of the 35S promoter to generate mC121-OmAGAL2, mC121-ΔSP-OmAGAL2, and mC121-SP, respectively. Transient expression was conducted by co-inoculation of *Agrobacterium tumefaciens* strain GV3101 cultures carrying each construct with those carrying the vector pMDC-At5g11420:pH-tdGFP as an apoplast marker (Stoddard and Rolland, 2019), and pDGB3alph2_35S:P19:Tnos (GB1203, Addgene #68214) in leaves of *Nicotiana benthamiana*. mCherry and green fluorescent protein (GFP) were excited at 555 and 488 nm, and observed in the range of 570–600 and 490–520 nm, respectively, using a LSM700 laser scanning confocal microscope (Carl Zeiss, Jena, Germany).

### Agrobacterium tumefaciens

GV3101 cultures carrying mC121-OmAGAL2, mC121-ΔSP-OmAGAL2, and mC121-SP, were used to inoculate Arabidopsis plants by floral inoculation and the T_2_ generations of transgenic plants were obtained. Five- to 8-day-old seedlings were treated with ClearSee solution (Kurihara *et al*, 2015), stained with Calcofluor White Stain (CWS, Merck) and observed using a LSM700 laser scanning confocal microscope. CWS was excited at 405 nm and observed in the range of 400–520 nm.

### Protein expression in tobacco BY-2 Cells

*Agrobacterium tumefaciens* GV3101 cultures carrying mC121-OmAGAL2, mC121-ΔSP-OmAGAL2, and mC121-SP were used to infect tobacco BY-2 cells.

Transgenic BY-2 cells were cultured in liquid Murashige and Skoog medium in a rotary shaker at 80 rpm at 25 °C. Cells and medium were separately collected by filtration 4 days after subculture. The culture medium was concentrated using Amicon Ultra-15 Centrifugal Filter Units, 10 kDa (Merck), desalted with PD 10 desalting columns (Merck) in extraction buffer (50 mM HEPES with 1mM EDTA, 1mM 2-ME, 0.1% (w/v) NaN_3_, and 0.1% (v/v) Protease Inhibitor Cocktail (Promega, Madison, MA, USA), pH 7.0) and used as apoplast protein fractions. The collected BY-2 cells were ground in liquid nitrogen with a mortar and pestle, and soluble proteins were extracted in the extraction buffer.

### Western-blot analysis

Proteins were separated on 10% (w/v) SDS-PAGE and electrically transferred to a PVDF membrane (Trans-Blot Turbo Mini 0.2 µm PVDF Transfer Packs, Bio-Rad, Hercules, CA, USA) using the Trans-Blot Turbo Transfer System (Bio-Rad) (1.3 A, 25 V, 10 min). The membrane was blocked with 5% (w/v) skim milk in TBST buffer (20 mM Tris-HCl, 150 mM NaCl, and 0.2% (w/v) Tween 20, pH 7.6) (blocking solution) for 1 h with gentle shaking. After washing with TBST buffer, the membrane was incubated with rabbit anti-mCherry polyclonal antibody (Proteintech Group, Rosemont, IL, USA) in blocking solution (1:6000) for 1.5 h. Multi Capture HRP (Fujifilm Wako Pure Chemical, Osaka, Japan) and ImmunoStar LD (Fujifilm Wako Pure Chemical) were used for detection of the antibody and chemiluminescence reactions, respectively, in accordance with the manufacture’s instruction. A luminescent image was captured using the Luminograph system (Atto, Tokyo, Japan).

### Accession numbers

*OmAGAL1* (LC636199), *OmAGAL2* (LC636200), *OmAGAL3* (LC636201), and comp71446_c0_seq1 (LC636330).

## Results

### Distribution of planteose in dry seeds of O. minor

Planteose is accumulated in the dry seeds of *O. minor* as a storage carbohydrate (Wakabayashi *et al.*, 2015). However, no tissue that accumulates planteose has been elucidated in any plant species. In the present study, MALDI–MSI was conducted to reveal the distribution of planteose in *O. minor* dry seeds. Two MS/MS fragment ions (m/z 527.16 [M+Na]^+^ > 365.11, 347.09) from planteose (Fig. 1A) were detected at 3 µm resolution (Fig. 1C, D). Images obtained for both fragment ions were almost identical, indicating that these fragment ions were generated from a single source, planteose. Strong signals were mainly detected from the seed coat, and some areas in the perisperm and endosperm, but not the embryo (Fig. 1B–D). This distribution pattern is in agreement with the role of planteose as a storage carbohydrate.

**Fig. 1.**
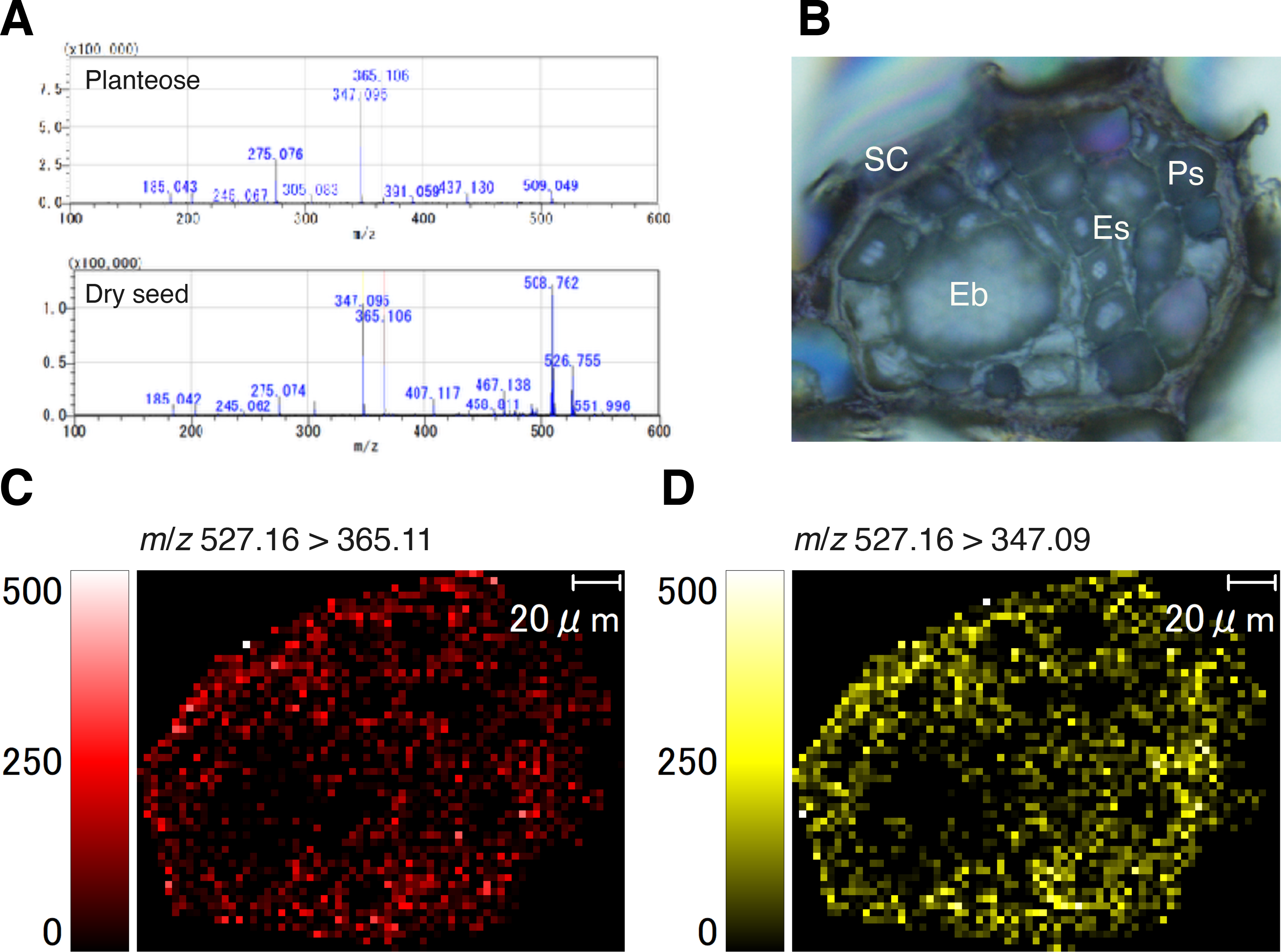
Mass imaging of planteose in dry seeds of *O*. *minor*. (A) MALDI–MS/MS spectra of purified planteose (upper) and *O*. *minor* dry seed (lower). 2,5-Dihydroxybenzoic acid was mixed with purified planteose or sprayed on a dry seed section as matrix, and *m*/*z* 527.16 [M+Na]^+^ was selected as a precursor ion. (B) Bright field image of an *O*. *minor* dry seed section. Eb, Embryo; Es, Endosperm, Ps, Perisperm; SC, Seed coat. Mass images of fragment ions *m*/*z* 365.11 (C) and 347.09 (D) from a precursor ion *m*/*z* 526.17.

### α-Galactosidase activity in germinating seeds of O. minor

Planteose was almost completely hydrolyzed by 5 days after *rac*-GR24 treatment (DAT) in *O. minor* (Wakabayashi *et al.*, 2015). Given that the first step of planteose metabolism is hydrolysis of an α-galactosyl linkage, AGAL activities in crude enzyme fractions prepared from germinating seeds of *O. minor* were measured under pH 5.0 or 7.0 using *p*-NP-α-Gal as a substrate. The AGAL activity at 1 DAT was lower than that after conditioning just before *rac*-GR24 treatment (0 DAT), then increased under both pH 5.0 and 7.0, although the activity under pH 5.0 was much higher than that under pH 7.0 (Fig. 2). These results indicated that acid AGAL was activated after perception of SL in germinating seeds of *O. minor*.

**Fig. 2.**
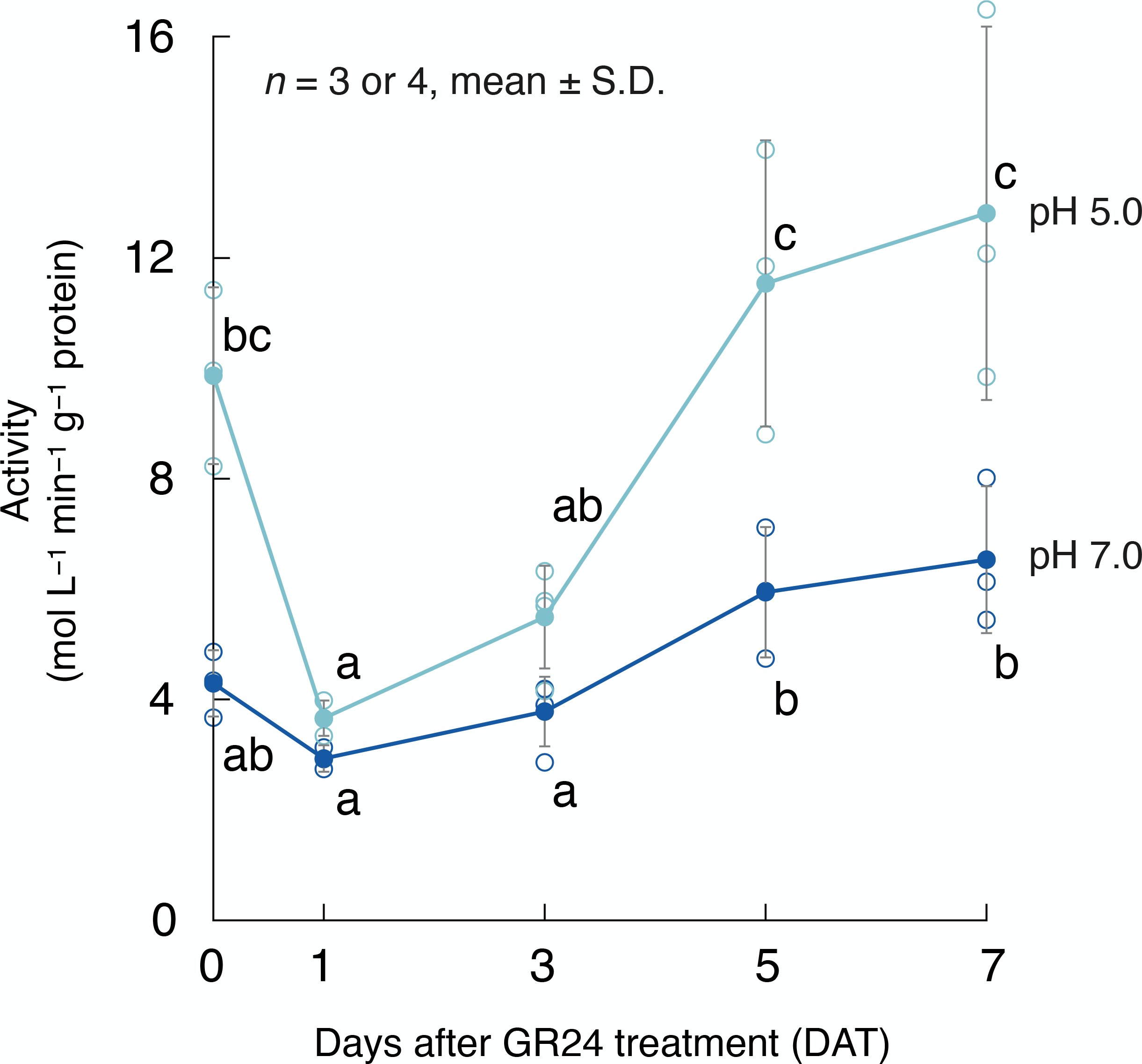
Changes in α-galactosidase activity in germinating seeds of *O*. *minor*. α-Galactosidase activity in the crude enzyme fraction prepared at different time points during germination was measured under pH 5.0 or 7.0 using *p*-nitrophenyl-α-D-galactopyranoside (1.0 mM) as a substrate. Values are means with standard deviations (*n* = 3 or 4). Different lowercase letters indicate significant difference under the same pH *(p* < 0.05, Tukey–Kramer test).

### In vivo visualization of α-galactosidase activity

MALD–MSI suggested that planteose accumulated in tissues other than the embryo, such as the endosperm, perisperm, and seed coat. We visualized AGAL activity in germinating seeds of *O*. *minor* using X-α-Gal. After X-α-Gal solution was applied to the germinating seeds, blue coloration, indicating hydrolysis of the α-galactosyl linkage in X-α-Gal, was observed in the seed coats near the micropyle where the radicle emerged at 3 and 5 DAT (Fig. 3A, B). Given that AGAL activity was detected near the embryo, we manually separated the embryo with the radicle from the endosperm and seed coat, and X-α-Gal solution was applied. The AGAL activity was confirmed within the seed coats, and was not detected on or in the embryo and radicle at 3 and 5 DAT (Fig. 3C, D). The localization of detected AGAL activity indicated that AGAL was involved in translocation of the storage carbohydrate into the embryo from the surrounding tissues.

**Fig. 3.**
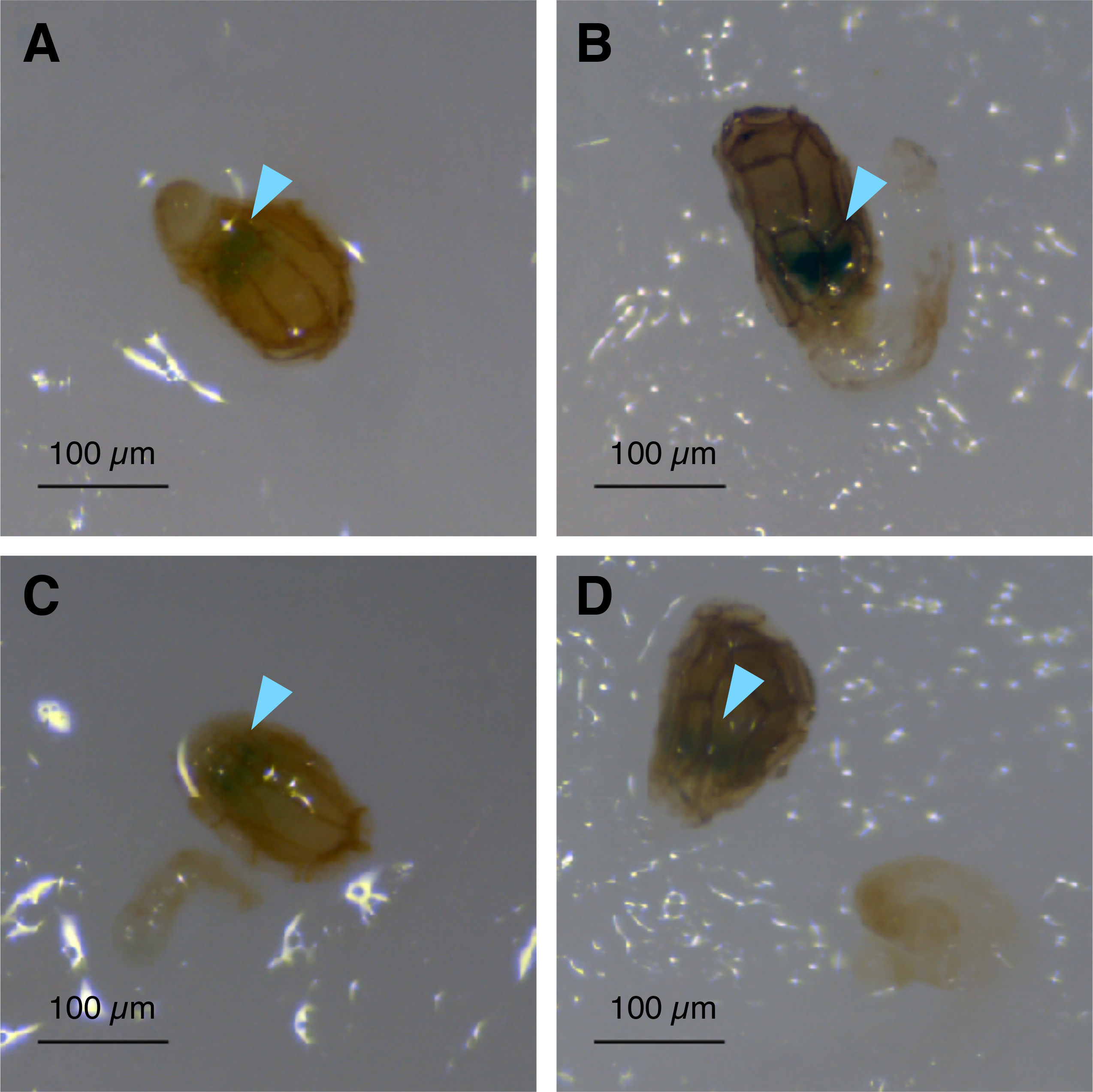
*In vivo* visualization of α-galactosidase activity in germinating seeds of *O*. *minor* with 5-bromo-4-chloro-3-indolyl-α-D-galactopyranoside (X-α-Gal). The seeds at (A) 3 days after *rac*-GR24 treatment (DAT) or (B) 5 DAT were treated with X-α-Gal solution (40 µg mL^−1^). The endosperm and seed coat were separated manually from the embryo with emerging radicles at (C) 3 DAT or (D) 5 DAT, and treated with X-α-Gal solution.

### Molecular cloning and characterization of α-galactosidase in O. minor

Transcriptomic data (Okazawa *et al.*, 2020) were surveyed for molecular cloning of AGAL genes in *O*. *minor*. Three candidate genes, comp64068_c0_seq7, comp35887_c0_seq1, and comp34431_c0_seq1, were annotated as AGAL-homologous genes and were named *OmAGAL1*, *OmAGAL2*, and *OmAGAL3*, respectively, based on the homology with *AtAGAL1* (At5g08380), *AtAGAL2* (At5g08370), and *AtAGAL3* (At3g56310) of Arabidopsis (Imaizumi *et al.*, 2017) (Supplemental Figs. S1, S2). Because the expression level of *OmAGAL2* was highest among these genes during germination (Supplemental Fig. 3), *OmAGAL2* was chosen for further analysis. A RT-qPCR assay confirmed that expression of *OmAGAL2* was induced after perception of *rac*-GR24 and peaked at 3–5 DAT (Fig. 4).

**Fig. 4.**
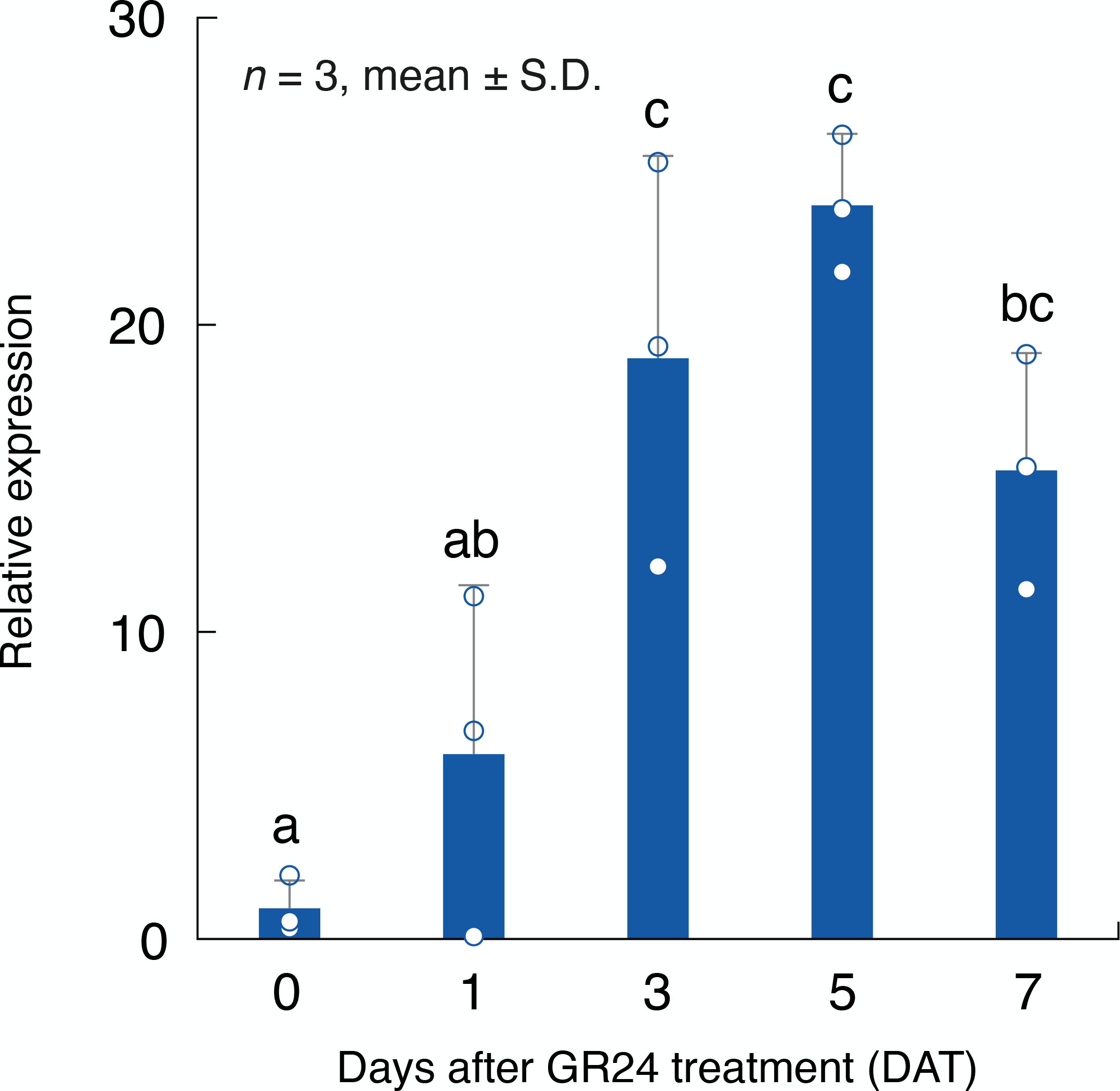
Expression of *OmAGAL2* during seed germination. *OmAGAL2* expression was quantified by RT-qPCR analysis. A constitutively expressed gene, comp71446_c0_seq1, was used as a reference gene for normalization of the expression level. Values are means with standard deviations (*n* = 3). Different lowercase letters indicate a significant difference *(p* < 0.05, Tukey–Kramer test).

The coding sequence of *OmAGAL2* was isolated from the transcriptome of germinating seeds of *O*. *minor*. The open reading frame was 1239 bp and encoded 412 amino acid residues with a predicted molecular mass of 45 kD. OmAGAL2 showed high homology with other AGALs in GH27, possessing a SP at its N-terminus, an α-galactosidase motif, and two conserved aspartate residues at active sites (Tapernoux-Lüthi *et al.*, 2004; Imaizumi *et al.*, 2017) (Supplemental Fig. 2). Phylogenetic analysis revealed that OmAGAL2 was placed in a clade including AtAGAL2 (Imaizumi *et al.*, 2017) and showed the highest homology with *S*. *asiatica* α-galactosidase (SaAGAL) (Supplemental Fig. 4). *Orobanche minor* and *S*. *asiatica* are both members of the Orobanchaceae.

OmAGAL2 lacking the SP (AA1–46), ΔSP-OmAGAL2 was heterologously expressed in *E*. *coli*. ΔSP-OmAGAL2 exhibited AGAL activity with optimal activity detected at pH 5.0−6.0 (Fig. 5A), indicating that OmAGAL2 is an acid AGAL active in acidic compartments. ΔSP-OmAGAL2 hydrolyzed planteose to Suc at pH 5.0 (Fig. 5B–D), whereas the activity was not confirmed at pH 7.0 (data not shown).

**Fig. 5.**
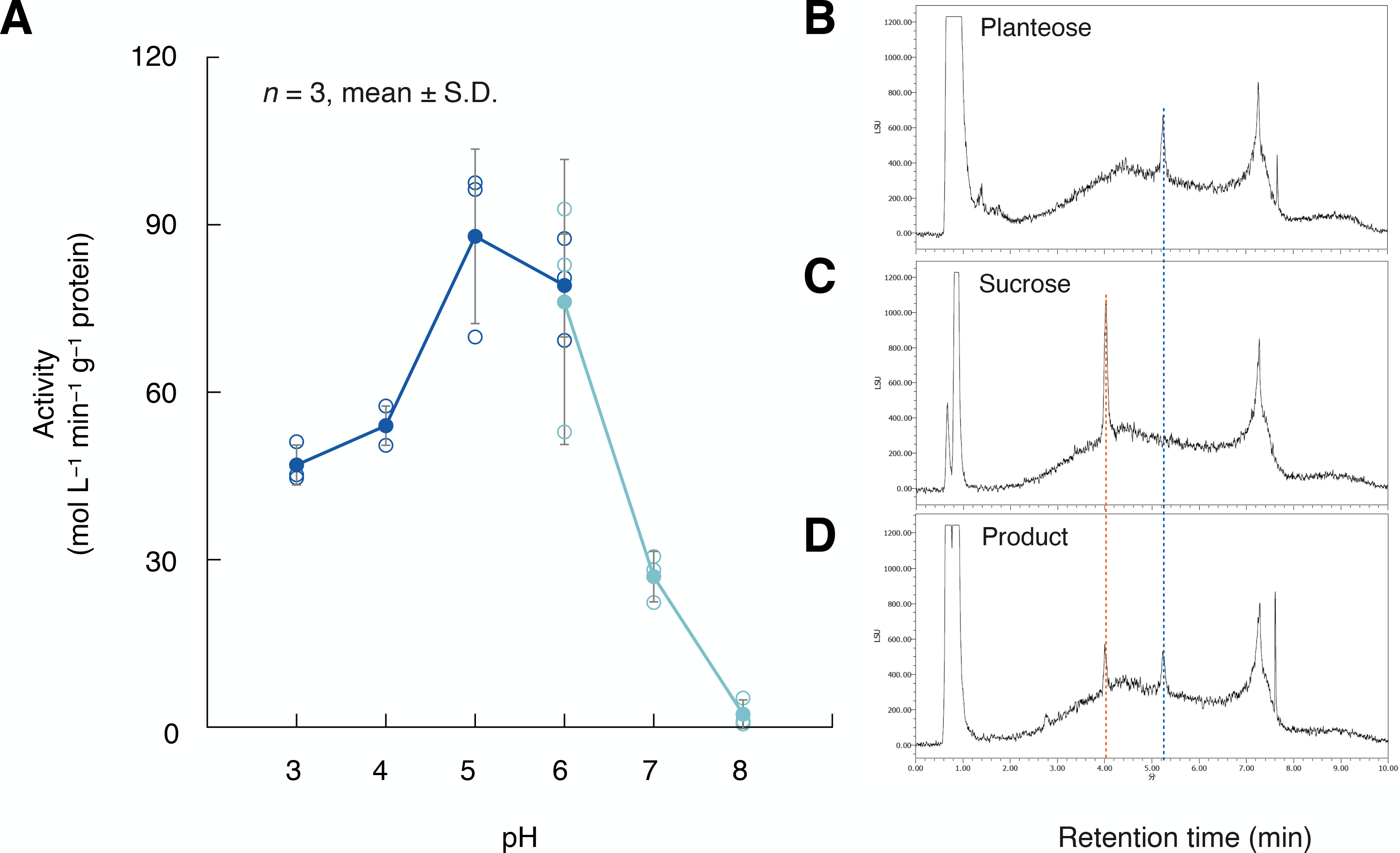
Enzymatic activity of ΔSP-OmAGAL2 expressed in *E*. *coli*. (A) Effect of pH on α-galactosidase activity of ΔSP-OmAGAL2. The activity was measured in 0.1 M citrate buffer (pH 3.0−6.0) or 0.1 M phosphate buffer (pH 6.0−8.0) at 37 °C for 30 min using *p*-NP-α-Gal (1.0 mM) as a substrate. Values are means with standard deviations (*n* = 3). (B–D) UPLC-ELSD chromatogram of purified planteose (B), authentic Suc (C), and product after incubation of planteose with ΔSP-OmAGAL2 under pH 5.0 (D). Blue and red dashed lines indicate the peak of planteose and Suc, respectively.

### Subcellular localization of OmAGAL2

The function of the N-terminal SP was evaluated using mCherry fusion constructs. OmAGAL2, ΔSP-OmAGAL2, and SP fused with mCherry at the C-terminus were transiently expressed in leaves of *N*. *benthamiana* together with an apoplast marker, At5g11420:pH-tdGFP (Stoddard and Rolland, 2019). mCherry fluorescence from OmAGAL2:mCherry and SP:mCherry overlapped with GFP fluorescence from At5g11420:pH-tdGFP, indicating that OmAGAL2 was secreted into apoplast as a result of the SP function (Fig. 6A-D, I-L). In contrast, ΔSP-OmAGAL2 and mCherry as a negative control were localized in the nucleus and cytoplasm (Fig. 6E-H, M-P).

**Fig. 6.**
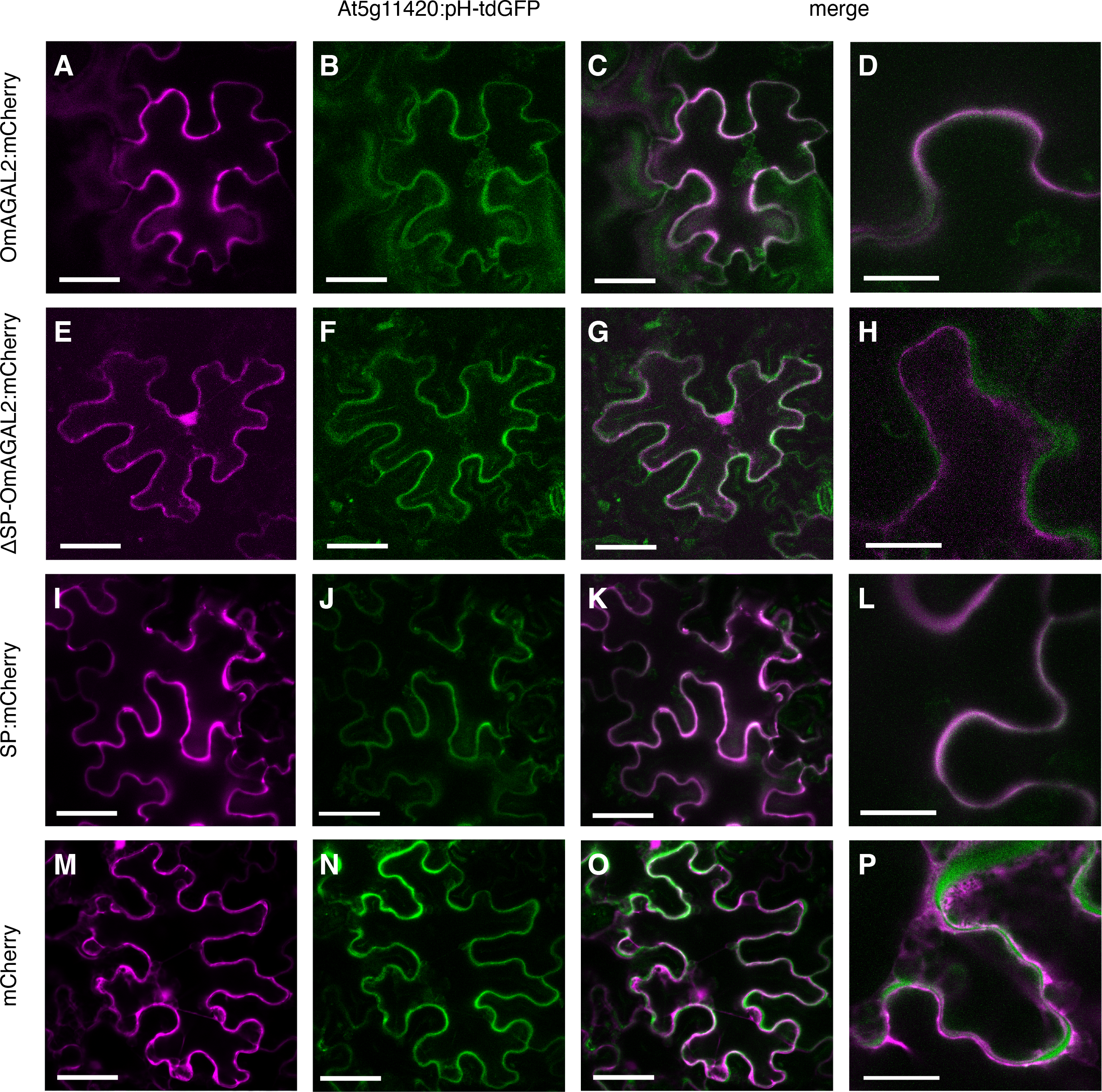
Localization of mCherry fusion proteins transiently expressed in leaves of *Nicotiana benthamiana*. (A) OmAGAL2:mCherry, (E) ΔSP-OmAGAL2:mCherry, (I) SP:mCherry, and (M) mCherey were co-expressed with the apoplast marker At5g11420:pH-tdGFP (B, F, J, N). (C, D, G, H, K, L, O, P) Merged images of fluorescence from mCherry and GFP. Scale bars: 50 µm (A–C, E–G, I–K, M–O) and 20 µm (D, H, L, P).

To confirm the transient assay results, Arabidopsis was transformed with the same constructs. Fluorescence from mCherry generated lattice-like patterns surrounding root cells, and strong fluorescence was observed in intercellular spaces in the case of OmAGAL2:mCherry and SP:mCherry (Fig. 7A-D, I-L). In contrast, fluorescence from ΔSP-OmAGAL2 was observed within the cells (Fig. 7E-H). Fluorescence from mCherry as a negative control was also detected within the cells and especially accumulated in the nucleus (Fig. 7M-P). The results of transient expression of the constructs in *N*. *benthamiana* and transgenic Arabidopsis were consistent and indicated that secretion of OmAGAL2 into the apoplast was dependent on SP function.

**Fig. 7.**
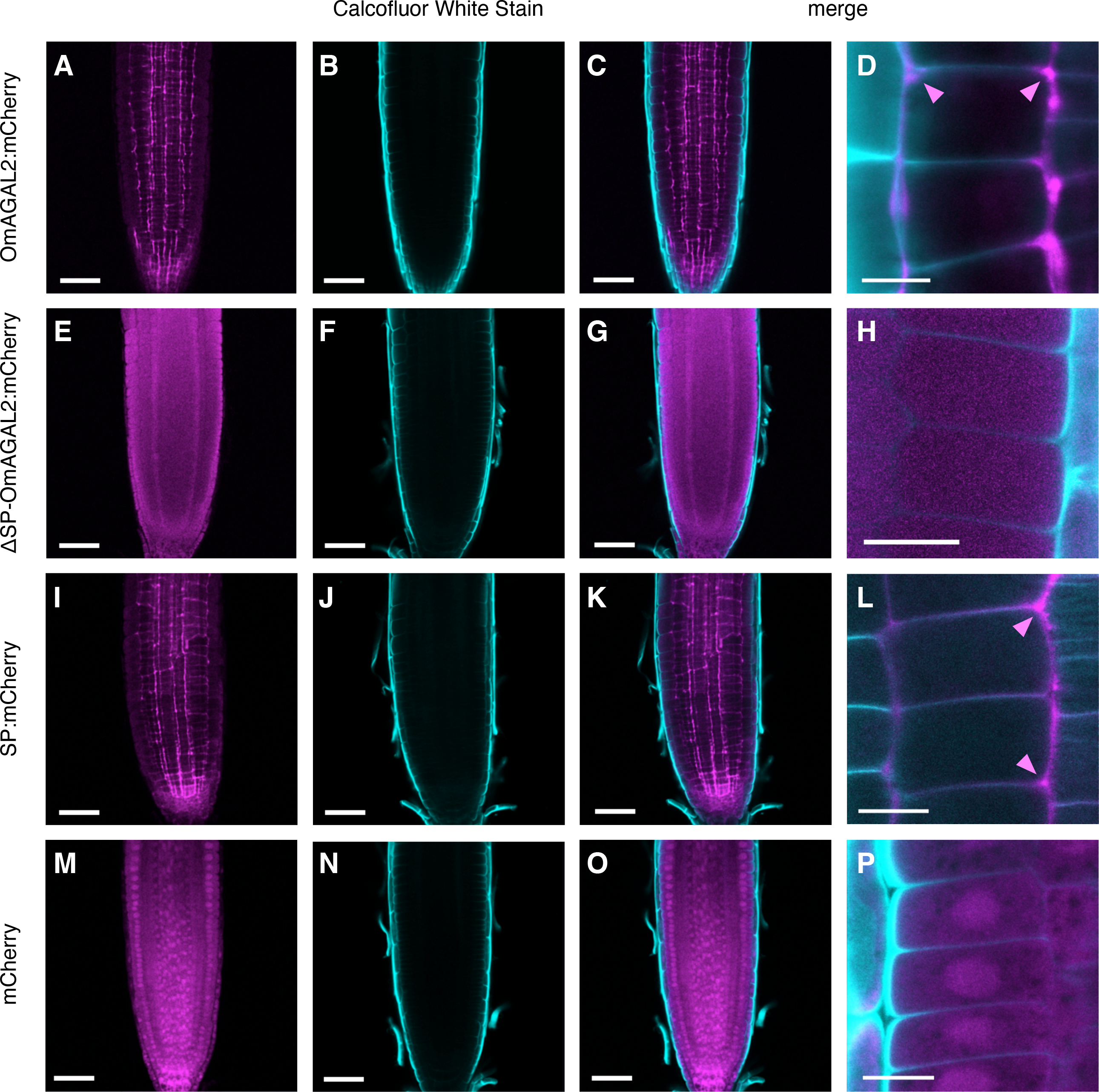
Localization of mCherry fusion proteins expressed in roots of transgenic Arabidopsis. (A) OmAGAL2:mCherry, (E) ΔSP-OmAGAL2:mCherry, (I) SP:mCherry, and (M) mCherry. Cell walls were stained with Calcofluor White Stain (CWS) (B, F, J, N). (C, D, G, H, K, L, O, P) Merged images of fluorescence from mCherry and CWS. Scale bars: 50 µm (A–C, E–G, I–K, M–O) and 20 µm (D, H, L, P).

Secretion of the proteins was evaluated by western blot analysis using tobacco BY-2 cells transformed with the same constructs. Anti-mCherry polyclonal antibody detected the corresponding proteins in protein extracts from transformed BY-2 cells (Fig. 8A), whereas secretion of SP-containing proteins was confirmed in the medium in which BY-2 cells were cultured (Fig 8B). Taken together, the present results demonstrated that the SP in OmAGAL2 functions as a signal for protein secretion into apoplast.

**Fig. 8.**
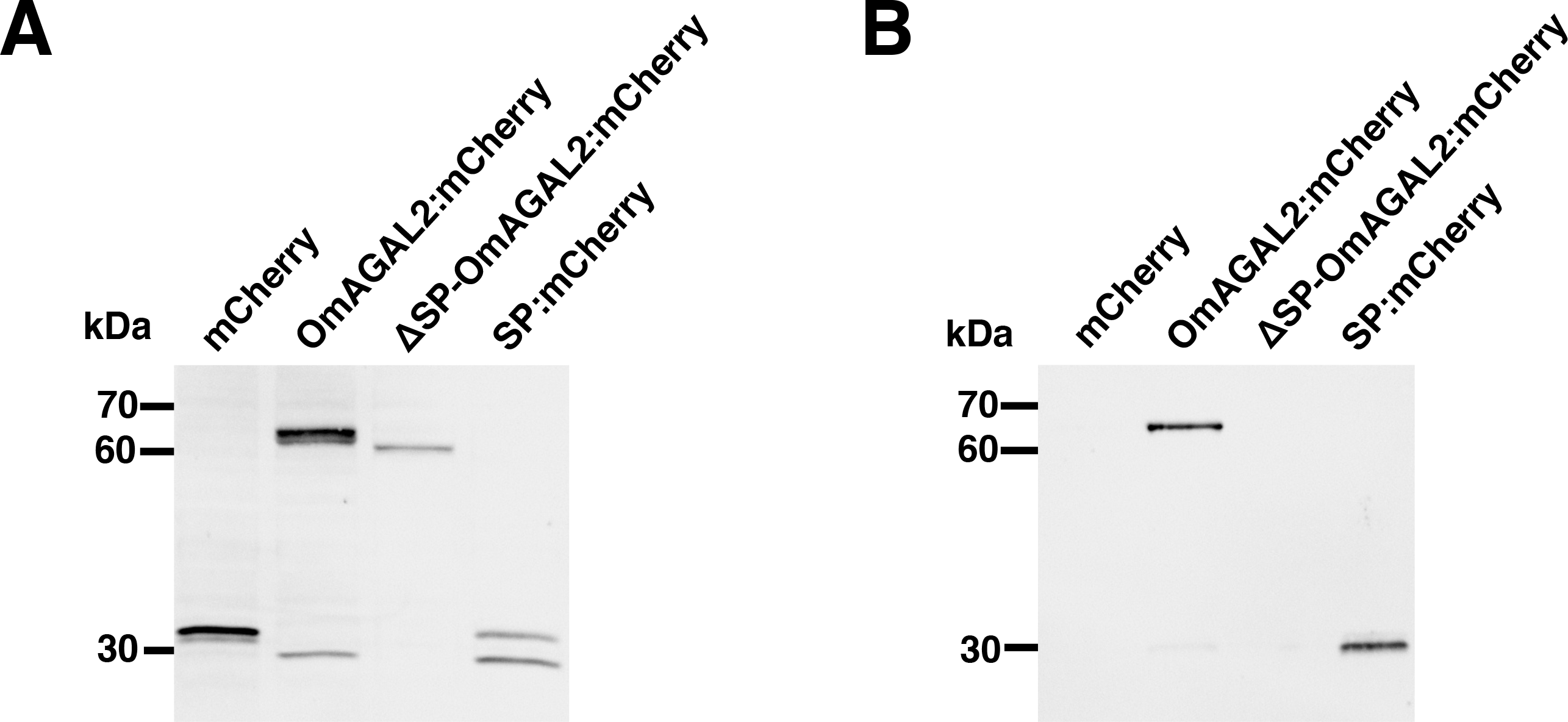
Secretion of proteins by the signal peptide (SP) from transgenic tobacco BY-2 cells. Western blot analysis of protein extraction from transgenic BY-2 cells (A) and medium in which BY-2 cells were cultured (B) with anti-mCherry polyclonal antibody.

## Discussion

### Planteose distribution in O. minor dry seeds

Planteose is contained in the seeds of several plant species, although its physiological role remains elusive. During seed germination of root parasitic weeds, planteose is rapidly hydrolyzed after perception of SLs, which is indicative of its role as a storage carbohydrate (Wakabasyhi *et al.*, 2015). In general, the endosperm stores storage carbohydrates and supplies nutrients to the embryo during germination, such as starch in cereals or lipids in Arabidopsis (Yan *et al.*, 2014). It is also considered that tissues surrounding the embryo, namely the endosperm, perisperm, and seed coat, play roles in nutrient supply in root parasitic weeds (Joel *et al.*, 2012; Joel and Bar, 2013). In the present study, we demonstrated that planteose is distributed in the endosperm, perisperm and seed coat in the dry seeds of *O. minor* (Fig. 1), which coincides with its role as a storage carbohydrate. Ultrastructural analysis of *P. aegyptiaca* seeds indicated that an endothelial cell layer beneath the seed coat surrounding the perisperm is filled with mucilage (Joel *et al.*, 2012). In chia seeds, planteose is a major oligosaccharide in the mucilage (Xing *et al.*, 2017). Accordingly, fragment ions detected in the seed coat might be from planteose accumulated in the endothelial mucilage (Fig. 1C, D). In addition, planteose was widely detected in the endosperm and perisperm, indicating that planteose is contained not only in the mucilage but also in compartments of the endosperm and perisperm. The use of MALDI–MSI to visualize carbohydrates in plant tissues, such as apple fruit (Horikawa *et al.*, 2019) and wheat stems (Robinson *et al.*, 2007), has proved to be a powerful technique to reveal carbohydrate distribution. In the current study, we visualized the distribution of the storage carbohydrate planteose in seeds of a root parasitic weed for the first time.

### α-Galactosidase expression during O. minor seed germination

Metabolic profiling previously revealed that the first step of planteose metabolism is hydrolysis of the α-galactosyl linkage (Wakabayashi *et al.*, 2015), although the corresponding enzymes have not been elucidated in any plant species. An enzymatic assay confirmed that AGAL activity increased with the progression of germination, especially under an acidic pH (Fig. 2). This result indicated that acid AGAL was activated in acidic compartments, i.e., the vacuole or apoplast. After X-α-Gal was applied to the germinating seeds to visualize AGAL activity, a blue coloration indicated that activity was localized near the micropyle (Fig. 3). Interestingly, if X-α-Gal was applied after separation of the embryo with the emerging radicle from the seed coat, blue coloration was confirmed only in the seed coat, suggesting that AGAL was expressed in tissues surrounding but not within the embryo (Fig. 3). This result was consistent with the MSI of planteose, which showed that planteose accumulated mainly outside of the embryo (Fig. 1). However, AGAL activity was detected specifically near the micropyle, whereas planteose was distributed in the non-embryonic tissues in the seed. Small perisperm cells surrounding the emerging radicle in the micropylar region are considered to supply nutrients to the embryo (Joel *et al.*, 2012). The current results indicated that AGAL is activated specifically in perisperm cells surrounding the embryo, thus suggesting the involvement of AGAL in nutrient transport. Whether planteose is translocated by a specific route or moves by passive diffusion to its hydrolytic site remains to be determined.

Characterization of AGALs in some plant species has been conducted in relation to metabolism of RFOs (Van den Ende, 2013) and galactomannan (Buckeridge *et al.*, 2000). Activation of AGAL during germination has been observed in plant species such as lettuce (*Lactuca sativa*) (Leung and Bewley, 1981), soybean (Guimarães *et al.*, 2001), coffee (*Coffea arabica*) (Marraccini *et al.*, 2005), and pea (Blöchl *et al.*, 2008). The AGAL activity in cell walls of the endosperm is increased during seed germination in date palm (Chandra Sekhar and DeMason, 1990). In tomato, acid AGAL activity is increased in the lateral and micropylar regions of the endosperm but not in the embryo during germination (Feurtado *et al.*, 2001). These observations indicate that AGALs are universally expressed during seed germination to mobilize stored carbohydrates to the embryo, although their substrate may differ among species, i.e. galactomannan, raffinose, or other oligosaccharides. Given that raffinose or galactomannan are not reported to be present in seeds of root parasitic weeds, and planteose is the only trisaccharide detected in our previous study (Wakabayashi *et al.*, 2015), planteose may be the substrate of AGAL in *O. minor* and other root parasitic weeds, as supported by the high similarity of OmAGAL2 and SaAGAL (Supplemental Fig. 4). Further characterization of polysaccharides in the cell wall of root parasitic weeds is a future challenge to elucidate the storage carbohydrates in their seeds.

### Characteristics of OmAGAL2

As a candidate planteose-hydrolytic enzyme, *OmAGAL2* was cloned and expressed in *E*. *coli* as ΔSP-OmAGAL2. OmAGAL2 belongs to the GH27 protein family, to which plant acid AGALs are assigned (Van den Ende, 2013). A putative SP at its N-terminus (AA1–32), and an AGAL motif and three aspartate residues required for recognition and hydrolysis of α-galactosyl saccharides (Tapernoux-Lüthi *et al.*, 2004; Imaizumi *et al.*, 2017) are conserved in OmAGAL2 (Supplemental Fig. 2). In comparison with biochemical studies, few molecular investigations have used recombinant AGALs from plants. Characterization of ArGGT1, an ortholog of AtAGAL1 in common bugle (*Ajuga reptans*), revealed its function as a raffinose oligosaccharide chain elongation enzyme rather than a hydrolytic enzyme (Tapernoux-Lüthi *et al.*, 2004) (Supplemental Fig. 3). AtAGAL2 and AtAGAL3 show activity toward RFOs, but RFOs are not expected to be their main substrate *in planta* (Imaizumi *et al.*, 2017). Involvement of AtAGAL3 in the hydrolysis of L-arabinopyranose residues in cell wall components has been proposed because of the enzyme’s β-L-arabinopyranosidase activity (Imaizumi *et al.*, 2017). Rice AGALs also hydrolyze RFOs and galactomannans, but their intrinsic substrates have not determined to date (Li *et al.*, 2007). As described herein, planteose is a possible substrate of AGALs and, in the current study, OmAGAL2 was demonstrated to hydrolyze planteose at pH 5.0. Moreover, OmAGAL2 was proven to be secreted to the apoplast through the function of the N-terminus SP. Accordingly, planteose is likely to be hydrolyzed in the acidic apoplast by OmAGAL2.

### Hypothetical model of planteose mobilization in germinating seeds of O. minor

From the present results, we propose a hypothetical model of planteose mobilization during seed germination of *O*. *minor*. Planteose is stored outside of the embryo in some compartments, such as the endothelial mucilage, endosperm, and perisperm. After perception of SLs, transcription of OmAGAL2 is up-regulated and the transcript level peaks at 5 DAT when planteose is completely hydrolyzed. The expression of OmAGAL2 might be restricted to tissues surrounding the embryo near the micropyle, as indicated by visualization of AGAL activity using X-α-Gal. Given that nutrients are mobilized from the endosperm and perisperm into the embryo, the localized AGAL activity emphasizes its involvement in the hydrolysis of storage carbohydrates. Sucrose released from planteose by OmAGAL2 may be directly translocated into embryonic cells through Suc transporters (SUTs). However, SUTs are unlikely to be involved in germination, because application of exogenous Suc does not induce the germination of NJ-treated *O*. *minor* seeds. However, NJ-treated seeds were germinated in response to application of exogenous Glc (Wakabayashi *et al.*, 2015). Therefore, Suc might be further hydrolyzed by cell wall invertases. Involvement of invertases in seed germination in *O*. *minor* (Wakbayashi *et al.*, 2015) and *P. ramosa* (Draie *et al.*, 2011) has been documented previously. Hexoses may be incorporated into embryonic cells through hexose transporters.

Previously, we observed that NJ suppressed seed germination of *O*. *minor* through inhibition of planteose metabolism (Wakabayashi *et al.*, 2015; Harada *et al.*, 2017; Okazawa *et al.*, 2020). The present results suggest that OmAGAL2 is a potential target for control of root parasitic weeds. The apoplastic localization of OmAGAL2 is a favorable characteristic for inhibitor design because consideration of the membrane permeability of inhibitors can be omitted. Screening of OmAGAL2 inhibitors is ongoing using recombinant proteins, and evaluation of the effects of the inhibitors on carbohydrate metabolism and germination in *O*. *minor* is the focus of an ensuing study.

### Supplementary data

The following supplementary data are available at *JXB* online.

Fig. S1. Alignments of partial sequences of OmAGAL1 with Arabidopsis α-galactosidase AtAGAL1, and OmAGAL3 with AtAGAL3.

Fig. S2. Multiple alignment of OmGAL2 with AtAGAL1, AtAGAL2, and AtAGAL3.

Fig. S3. Expression of OmAGAL1, OmGAL2, and OmAGAL3 in germinating seeds of *O*. *minor*.

Fig. S4. Phylogenetic tree for plant AGALs.

## Acknowledgements

This work was partly supported by the Japan Society for the Promotion of Science (JSPS) KAKENHI (grant no. JP20H02924 to A.O. and D.O., JP20KK0130 to A.O. and Y.S.) and by Japan Science and Technology Agency (JST)/Japan International Cooperation Agency (JICA) Science and Technology Research Partnership for Sustainable Development (SATREPS) (grant no. JPMJSA1607 to A.O. and Y.S.). The authors would like to thank Dr. Vivien Rolland for providing pMDC-At5g11420:pH-tdGFP, and Yumi Mori, Maki Hazama, and Hiromi Saito for technical assistance. We thank Robert McKenzie, PhD, from Edanz Group, for editing a draft of this manuscript.

## Author contributions

A.O conceived the study, designed and supervised the research, analyzed the data, and wrote the article with the support of all authors; A.B., T.T., H.O., M.O. and T.N. performed most of the experiments and analyzed the data; S.S. conducted mass imaging; T.O, Y.S., and D.O. supervised the research.

## Data availability

All data supporting the findings of this study are available within the paper and within the supplementary data published online

## References

1. Akiyama K, Matsuzaki K, Hayashi H. 2005. Plant sesquiterpenes induce hyphal branching in arbuscular mycorrhizal fungi. Nature 435, 824–827.

2. Al-Babili S, Bouwmeester HJ. 2015. Strigolactones, a novel carotenoid-derived plant hormone. Annual Review of Plant Biology 66, 161–186.

3. Arite T, Umehara M, Ishikawa S, Hanada A, Maekawa M, Yamaguchi S, Kyozuka J. 2009. *D14*, a strigolactone-insensitive mutant of rice, shows an accelerated outgrowth of tillers. Plant and Cell Physiology 50, 1416–1424.

4. Blöchl A, Peterbauer T, Hofmann J, Richter A. 2008. Enzymatic breakdown of raffinose oligosaccharides in pea seeds. Planta 228, 99–110.

5. Bouwmeester HJ, Fonne-Pfister R, Screpanti C, De Mesmaeker A. 2019. Strigolactones: plant hormonse with promising features. Angewante Chemie International Edition 58, 12778–12786.

6. Bouwmeester H, Li C, Thiombiano B, Rahimi M, Dong L. 2021. Adaptation of the parasitic plant lifecyle: Germination is controlled by essential host signaling molecules. Plant Physiology 185, 1292–1308.

7. Brun G, Braem L, Thoiron S, Gevaert K, Goormachtig S, Delavault P. 2018. Seed germination in parasitic plants: what insights can we expect from strigolactone research? Journal of Experimental Botany 69, 2265–2280.

8. Buckeridge M, dos Santos HP, Tiné MAS. 2000. Mobilisation of storage cell wall polysaccharides in seeds. Plant Physiology and Biochemistry 38, 141–156.

9. Bürger M, Chory J. 2020. The many models of strigolactone signaling. Trends in Plant Science 25, 395–405.

10. Chandra Sekhar KN, DeMason DA. 1990. Identification and immunocytochemical localization of α-galactosidase in resting and germinated date palm (*Phoenix dactylfera* L.) seeds. Planta 181, 53–61.

11. Conn CE, Bythell-Douglas R, Neumann D, Yoshida S, Whittington B, Westwood JH, Shirasu K, Bond CS, Dyer KA, Nelson DC. 2015. Convergent evolution of strigolactone perception enabled host detection in parasitic plants. Science 349, 540–543.

12. Conn CE, Nelson DC. 2016. Evidence that KARRIKIN-INSENSITIVE2 (KAI2) receptors may perceive an unknown signal that is not karrikin or strigolactone. Frontiers in Plant Science 6, 1219.

13. Dey PM. 1980. Biosynthesis of planteose in *Sesamum indicum*. FEBS Letters 114, 153–156.

14. Draie R, Péron T, Pouvreau J-B, Véronési C, Jégou S, Delavault P, Thoiron S, Simier P. 2011. Invertases involved in the development of the parasitic plant *Phelipanche ramosa*: characterization of the dominant soluble acid isoform, PrSAI1. Molecular Plant Pathology 12, 638–652.

15. Fernández-Aparicio M, Delavault P, Timko MP. 2020. Management of infection by parasitic weeds: a review. Plants 9, 1184.

16. Feurtado JA, Banik M, Bewley D. 2001. The cloning and characterization of α-galactosidase present during and following germination of tomato (*Lycopersicon esculentum* Mill.) seed. Journal of Experimental Botany 52, 1239–1249.

17. Fialho LS, Guimarães VM, Callegari CM, Reis AP, Barbosa DS, Borges EEL, Moreira MA, Rezende ST. 2008. Characterization and biotechnological application of an acid α-galactosidase from *Tachigali multijuga* Benth. Seeds. Phytochemistry 69, 2579–2585.

18. French D. 1955. Isolation and identification of planteose from tobacco seeds. Journal of American Chemical Society 77, 1024–1025.

19. French D, Youngquist RW, Lee A. 1959. Isolation and crystallization of planteose from mint seeds. Archives of Biochemistry and Biophysics 85, 471–473.

20. Gomez-Roldan V, Fermas S, Brewer PB, et al. 2008. Strigolactone inhibition of shoot branching. Nature 455, 189–194.

21. Guimarães VM, de Rezende ST, Moreira MA, de Barros EG, Felix CR. 2001. Characterization of α-galactosidase from germinating soybean seed and their use for hydrolysis of oligosaccharides. Phytochemistry 58, 67–73.

22. Hamiaux C, Drummond RSM, Janssen BJ, Ledger SE, Cooney JM, Newcomb RD, α β hydrolase likely to be involved in the perception of the plant branching hormone, strigolactone. Current Biology 22, 2032–2036.

23. Harada K, Kurono Y, Nagasawa S, et al. 2017. Enhanced production of nojirimycin *via Streptomyces ficellus* cultivation using marine broth and inhibitory activity of the culture for seeds of parasitic weeds. Journal of Pesticide Science 42, 166–171.

24. Horikawa K, Hirama T, Shimura H, Jitsuyama Y, Suzuki T. 2019. Visualization of soluble carbohydrate distribution in apple fruit flesh utilizing MALDI-TOF MS imaging. Plant Science 278, 107–112.

25. Hughes SG, Overbeeke N, Robinson S, Pollock K, Smeets FLM. 1988. Messenger RNA from isolated aleurone cells directs the synthesis of an alpha-galactosidase found in the endosperm during germination of guar (*Cyampsis tetragonaloba*) seed. Plant Molecular Biology 11, 783–789.

26. Imaizumi C, Tomatsu H, Kitazawa K, Yoshimi Y, Shibano S, Kikuchi K, Yamaguchi M, Kaneko S, Tsumuraya Y, Kotake T. 2017. Heterologous expression and characterization of an Arabidopsis β-L-arabinopyranosidase and α-D-galactosidases acting on β-L-arabinopyranosyl residues. Journal of Experimental Botany 68, 4651–4661.

27. Joel DM, Bar H. 2013. The seed and the seedling. In: D Joel, J Gressel, L Musselman, eds, Parasitic Orobanchaceae, Springer, 147–166.

28. Joel DM, Bar H, Mayer AM, Plakhine D, Ziadne H, Westwood JH, Welbaum GE. 2012. Seed ultrastructure and water absorption pathway of the root-parasitic plant *Phelipanche aegyptiaca* (Orobanchaceae). Annal of Botany 109, 181–195.

29. Khosla A, Nelson DC. 2016. Strigolactones, super hormones in the fight against *Striga*. Current Opinion in Plant Biology 33, 57–63.

30. Kuo TM, VanMiddlesworth JF, Wolf WJ. (1988) Content of raffinose oligosaccharides and sucrose in various plant seeds. Journal of Agricultural and Food Chemistry 36, 32–36.

31. Kurihara D, Mizuta Y, Sato Y, Higashiyama T. 2015. ClearSee: a rapid optical clearing reagent for whole-plant fluorescence imaging. Development 142, 4168–4179.

32. Lanfranco L, Fiorilli V, Venice F, Bonfante P. 2018. Strigolactones cross the kingdoms: plants, fungi, and bacteria in the arbuscular mycorrhizal symbiosis. Journal of Experimental Botany 69, 2175–2188.

33. Leung DWM, Bewley JD. 1981. Red-light- and gibberellic-acid-enhanced α-galactosidase activity in germinating seeds, cv. Grand Rapids. Planta 152, 436–441.

34. Li S, Kim W-D, Kaneko S, Prema PA, Nakajima M, Kobayashi H. 2007. Expression of rice (*Oryza sativa* L. var. Nipponbare) α-galactosidase genes in Escherichia coli and characterization. Bioscience, Biotechnology, and Biochemistry 71, 520–526.

35. López-Ráez JA. 2015. How drought and salinity affect arbuscular mycorrhizal symbiosis and strigolactone biosynthesis? Planta 243, 1375–1385.

36. Marraccini P, Rogers WJ, Caillet V, Deshayes A, Granato D, Lausanne F, Lechat S, Pridmore D, Pétiard V. 2005, Biochemical and molecular characterization of α-D-galactosidase from coffee beans. Plant Physiology and Biochemistry 43, 909–920.

37. Nakamura H, Xue Y-L, Miyakawa T, et al. 2013. Molecular mechanism of strigolactone perception by DWARF14. Nature Communications 4, 2613.

38. Okazawa A, Kusunose T, Ono E, Kim HJ, Satake H, Shimizu B, Mizutani M, Seki H, Muranaka T. 2014. Glucosyltransferase activity of *Arabidopsis* UGT71C1 towards pinoresinol and lariciresinol. Plant Biotechnology 31, 561–566.

39. Okazawa A, Wakabayashi T, Muranaka T, Sugimoto Y, Ohta D. 2020. The effect of nojirimycin on the transcriptome of germinating *Orobanche minor* seeds. Journal of Pesticide Science 45, 230 237.

40. Parker C. 2013. The parasitic weeds of the Orobanchaceae. In: Joel D, Gressel J, Musselman L, eds. Parasitic Orobanchaceae, Springer, 313–344.

41. Robinson A, Warburton K, Seymour M, Clench M, Thomas-Oaters J. 2007, Localization of water-soluble carbohydrates in wheat stems using imaging matrix-assisted laser desorption ionization mass spectrometry. New Phytologist 173, 438–444.

42. Seto Y, Yasui R, Kameoka H, et al. 2019. Strigolactone perception and deactivation by a hydrolase receptor DWARF14. Nature Communications 10, 191.

43. Stoddard A, Rolland V. 2019. I see the light! Fluorescent proteins suitable for cell wall/apoplast targeting in *Nicotiana benthamiana* leaves. Plant Direct 3, e00112.

44. Tapernoux-Lüthi EM, Böhm A, Keller F. 2004. Cloning, functional expression, and characterization of the raffinose oligosaccharide chain elongation enzyme, galactan:galactan galactosyltransferase, from common bugle leaves. Plant Physiology 134, 1377–1387.

45. Tsuchiya Y, Yoshimura M, Sato Y, et al. 2015. Probing strigolactone receptors in *Striga hermonthica* with fluorescence. Science 349, 864–868.

46. Umehara M, Hanada A, Yoshida S, et al. 2008. Inhibition of shoot branching by new terpenoid plant hormones. Nature 455, 195–200.

47. Uraguchi D, Kuwata K, Hijikata Y, et al. 2018. A femtomolar-range suicide germination stimulant for the parasitic plant *Striga hermonthica*. Science 362, 1301–1305.

48. Van den Ende W. 2013. Multifunctional fructans and raffinose family oligosaccharides. Frontiers in Plant Science 4, 247.

49. Xing X, Hsieh YSY, Yap K, Ang ME, Lahnstein J, Tucker MR, Burton RA, Bulone V. 2017. Isolation and structural elucidation by 2D NMR of planteose, a major oligosaccharide in the mucilage of chia (*Salvia hispanica* L.) seeds. Carbohydrate Polymers 175, 231–240.

50. Yan D, Duermeyer L, Leoveanu C, Nambara E. 2014. The functions of the endosperm during seed germination. Plant and Cell Physiology 55, 1521−1523.

51. Yoneyama K. 2020. Recent progress in the chemistry and biochemistry of strigolactones. Journal of Pesticide Science 45, 45–53.

52. Yoshida S, Kim S, Wafula EK, et al. 2019. Genome sequence of *Striga asiatica* provides insight into the evolution of plant parasitism. Current Biology 29, 3041–3052.

53. Wakabayashi T, Joseph B, Yasumoto S, et al. 2015. Planteose as a storage carbohydrate required for early stage of germination of *Orobanche minor* and its metabolism as a possible target for selective control. Journal of Experimental Botany 66, 3085–3097.

54. Waters MT, Gutjahr C, Bennett T, Nelson DC. 2017. Strigolactone signaling and evolution. Annual Review of Plant Biology 68, 291–322.

